# Analyzing the neuroglial brainstem circuits for respiratory chemosensitivity in freely moving mice

**DOI:** 10.1101/492041

**Authors:** Amol Bhandare, Joseph van de Wiel, Reno Roberts, Ingke Braren, Robert Huckstepp, Nicholas Dale

**Affiliations:** School of Life Sciences, University of Warwick, Coventry, UK; University Medical Center Eppendorf, Vector Facility, Inst. for Exp. Pharmacology and Toxikology, N30, Room 09, Martinistr. 52, 20246 Hamburg, Germany

## Abstract

The regulated excretion of CO_^2^_ during breathing is a key life-preserving homeostatic mechanism. In the rostral medulla oblongata, neurons in two nuclei -the retrotrapezoid nucleus (RTN) and the rostral medullary Raphe -have been proposed as central CO_2_ chemosensors that mediate adaptive changes in breathing. One such adaptive change is active expiration, thought to be controlled by the lateral parafacial region (pF_L_). Here we use promoter-driven expression of GCaMP6 and head-mounted mini-microscopes to image the Ca^2+^ activity in neurons and glia of RTN, Raphe and pF_L_ in awake adult mice during inspiration of elevated CO_2_ to discriminate their roles in the adaptive response to hypercapnia. While there were multiple types of neuronal response to hypercapnia in both the RTN and the Raphe, a substantial proportion of neurons in the Raphe encoded the inspired level of CO_2_. By contrast, such responses were very rare in the RTN and the responses of RTN neurons suggest that this nucleus plays an integrative rather than a direct sensing role in the chemosensory reflex. pF_L_ neurons were reliably activated by hypercapnia and their activity preceded active expiration suggesting a key causal link to this aspect of breathing. Our analysis considerably revises understanding of chemosensory control in the adult awake mouse and paves the way to understanding how breathing is coordinated with complex non-ventilatory behaviours such as swallowing and vocalisation.

## Introduction

The precise control of breathing is fundamental to the survival of all terrestrial vertebrates. Breathing fulfils two essential functions: provision of oxygen to support metabolism; and removal of the metabolic by-product, CO_2_. Rapid and regulated removal of CO_2_ is essential because its over-accumulation in blood will result in death from the consequent drop in pH. The CO_2_-dependent regulation of breathing is mediated by chemosensitive neurons and glia in the medulla oblongata ^1-3^. In the rostral region of the medulla oblongata evidence favours the involvement of two neuronal populations: the Phox2B+ neurons of the retrotrapezoid nucleus (RTN) ^4-8^ and the serotonergic neurons of the rostral medullary raphe ^9-12^. Assessment of the relative roles of the RTN and Raphe has been hampered by the inability to record the activity of cells in these nuclei in awake freely behaving adult rodents during a hypercapnic challenge. Instead neuronal recordings have been made from young <14d or adult rodents under anaesthesia. Both of these methods have drawbacks -the chemosensory control of breathing matures postnatally; and anaesthesia is known to depress the activity of respiratory neurons and reconfigure the circuit.

During resting eupneic breathing, there is little active expiration instead the expiratory step involves elastic rebound of the respiratory muscles to push air out of the lungs. However when the intensity of breathing is increased e.g. during exercise or hypercapnia, active expiration (recruitment of abdominal muscles to push air out of the lungs) occurs. Mounting evidence suggests that the lateral parafacial nucleus (pF_L_) may contain the expiratory oscillator and that neurons in this nucleus should be recruited to evoke active expiration ^13-15^. Nevertheless the key step of recording activity of these neurons in awake behaving animals and linking it to active expiration has not been achieved.

Recently, deep brain imaging techniques have been developed to allow recording of activity of defined neuronal populations in awake, freely-moving animals ^16-18^. These methods requires: the expression of genetically encoded Ca^2+^ indicators such as GCaMP6 in the relevant neurons; the implantation of gradient refractive index (GRIN) lenses at the correct stereotaxic position; and a head-mounted mini-epifluorescence microscope to enable image acquisition during the free behaviour of the mouse. These methods have allowed imaging in deep brain structures such as the hypothalamus. We have now adapted them to enable recording of defined neuronal and glial populations in the brainstem of mice to analyze the chemosensory control of breathing and the contribution of pF_L_ to active expiration.

## Results

### Chemosensory responses in the RTN

Neurons of the RTN were targeted with a synapsin-GCaMP6s AAV (Fig 1A-C), a necessary step to enable their imaging during free behaviour via a mini-headmounted microscope (Fig 1B). The localization of successful and accurate transduction was confirmed posthoc by using choline acetyltransferase (ChAT) staining to define the facial nucleus and the location of the transduced cells (Fig 1D), with ∼ 7% of transduced neurons in the optical pathway of the microscope co-labelled with ChAT (Fig 1E).

**Figure 1:**
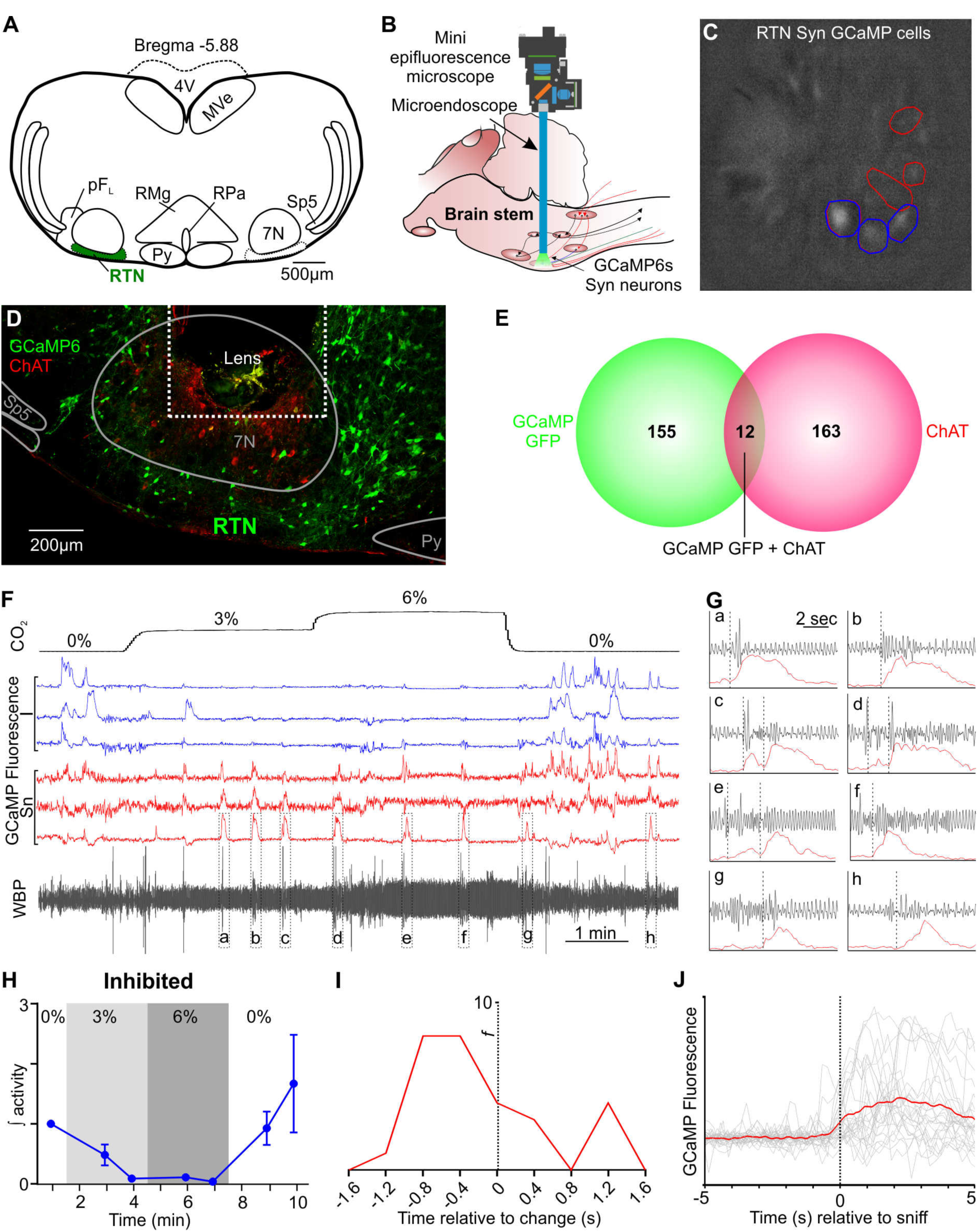
Sniff-coding and inhibited RTN neuronal reponses to elevated CO_2_ in awake mice. (A) AAV9-Syn-GCaMP6s injection into the RTN. (B) Representation of GRIN lens (microendoscope), baseplate, and mini epifluorescence camera placement for recording of brainstem nuclei. (C) GCaMP6s fluorescence of transduced RTN neurons in freely behaving mice. Individual regions of interest (ROIs) drawn around CO_2_-inhibited (blue) and sniff coactivated (red) neurons. (D) Micrograph of lens placement and viral transduction of neurons (green) relative to the facial nucleus (ChAT+ neurons - red). (E) Venn diagram of cell counts of GCaMP6s transduced (green), and ChAT+ (red), neurons under the GRIN lens. (F) Recording of mouse whole body plethysmography (WBP) in response to hypercapnia time-matched with Inscopix recorded GCaMP6 signals. CO_2_-Inhibited (blue) and sniff-coding (red) calcium traces correspond to the neurons shown in panel C. (G) Expanded recordings from (F), the start of the Ca^2+^ transient is shown by the dotted line. (H) Changes in activity (F/F_0_) of CO_2_-inhibited neurons. (I) Sniff-correlation histogram and (J) Spike triggered average (red line) of all Ca^2+^ events (grey lines) temporally correlated to the beginning of sniff activity (dotted verticle line). Abreviations: 7N, facial motor nucleus; Py, pyramidal tract; MVe, medial vestibular nucleus; sp5, spinal trigeminal nucleus: RTN, retrotrapezoid nucleus; RMg, raphe magnus; RPa, raphe pallidus; pF_L_, parafacial lateral region.

We recorded 143 RTN neurons in six mice. Surprisingly, the neuronal responses to the hypercapnic challenge were highly variable. To assist in classification, we subdivided neurons into 6 functional types according to how they responded to CO_2_: Excited - transient (E_T_), 9/143 showed a brief elevation in Ca^2+^ in response to a change in 3% inspired CO_2_; Excited - sustained (E_S_), 14/143 exhibited a sustained response to hypercapnia but which did not encode the degree of hypercapnia; Excited - graded (E_G_), 1/143 showed a graded response that encoded the level of inspired CO_2_; Excited - rebound (E_R_), 6/143 showed an elevated Ca^2+^ activity following the end of the hypercapnic episode; Inhibited (I), 9/143 were inhibited during the period of hypercapnia; and non-coding, having no discernable activity that related to the hypercapnic stimulus. In addition to these 6 categories, there was a seventh pattern of activity: sniff-coding (Sn), 3/143 showed elevated Ca^2+^ signals correlated with sniffing. Figure 1F shows examples of inhibited neurons and sniff-coding neurons recorded in the same field of view (Fig 1C and Supplementary Movie 1). Interestingly the different functional types of neuron are clustered adjacent to each other (Fig 1C). Ca^2+^ activity in the inhibited neurons was greatly reduced during both 3 and 6% inspired CO_2_ (Fig 1H) and often exhibited rebound following the end of the hypercapnic episode (Fig 1F and Supplementary Movie 1, blue traces). In sniff-coding neurons, Ca^2+^ activity coincided with perturbations of the plethysmography traces (increase in respiratory frequency, increase of tidal volume, presence of expiration, Fig 1G and Supplementary Movie 1, red traces). Analysis of the onset of the Ca^2+^ signal relative to the start of the sniff (Supplementary Fig 1), showed that, on average, activity in these neurons preceded the sniff by 0.4-0.8 s (Fig 1I). The temporal correlation between the sniff-coding neurons and the sniff was further confirmed by averaging the Ca^2+^ recordings from multiple sniffs (Fig 1J). Support for our evidence of sniff-coding neurons in the RTN comes from the observation that stimulation of sniffing leads to *c-Fos* expression in the retrofacial area of the cat (equivalent to the RTN) ^19^.

There were four classes of neurons that were excited by CO_2_ and indicated an aspect of the hypercapnic stimulus (Fig 2): E_T_ (the start of hypercapnia), E_S_ (the presence of hypercapnia), E_G_ (the presence and magnitude of the hypercapnic stimulus) and E_R_ (the post-hypercapnic period). The E_T_ responding neurons essentially encoded the beginning of hypercapnia (Fig 2A and Supplementary Movie 2, orange traces; Fig 2D). They preferentially responded to 3% inspired CO_2_ but could have some low level activity throughout the remained of the hypercapnic episode. E_S_ responding neurons showed elevated Ca^2+^ activity throughout the entire hypercapnic episode, but this did not encode the level of inspired CO_2_ (Fig 2A and Supplementary Movie 2, pink traces; Fig 2E). We observed only one E_G_ neuron in which the Ca^2+^ activity encoded the level of the CO_2_ stimulus (Fig 2B and Supplementary Movie 3; Fig 2F). The final class of responding neuron was E_R_. In these neurons there was sometimes elevated Ca^2+^ activity during the hypercapnic stimulus, but they were also characterised by a increase in activity following cessation of the hypercapnic episode (Fig 2C and Supplementary Movie 4, brown traces; Fig 2G).

**Figure 2:**
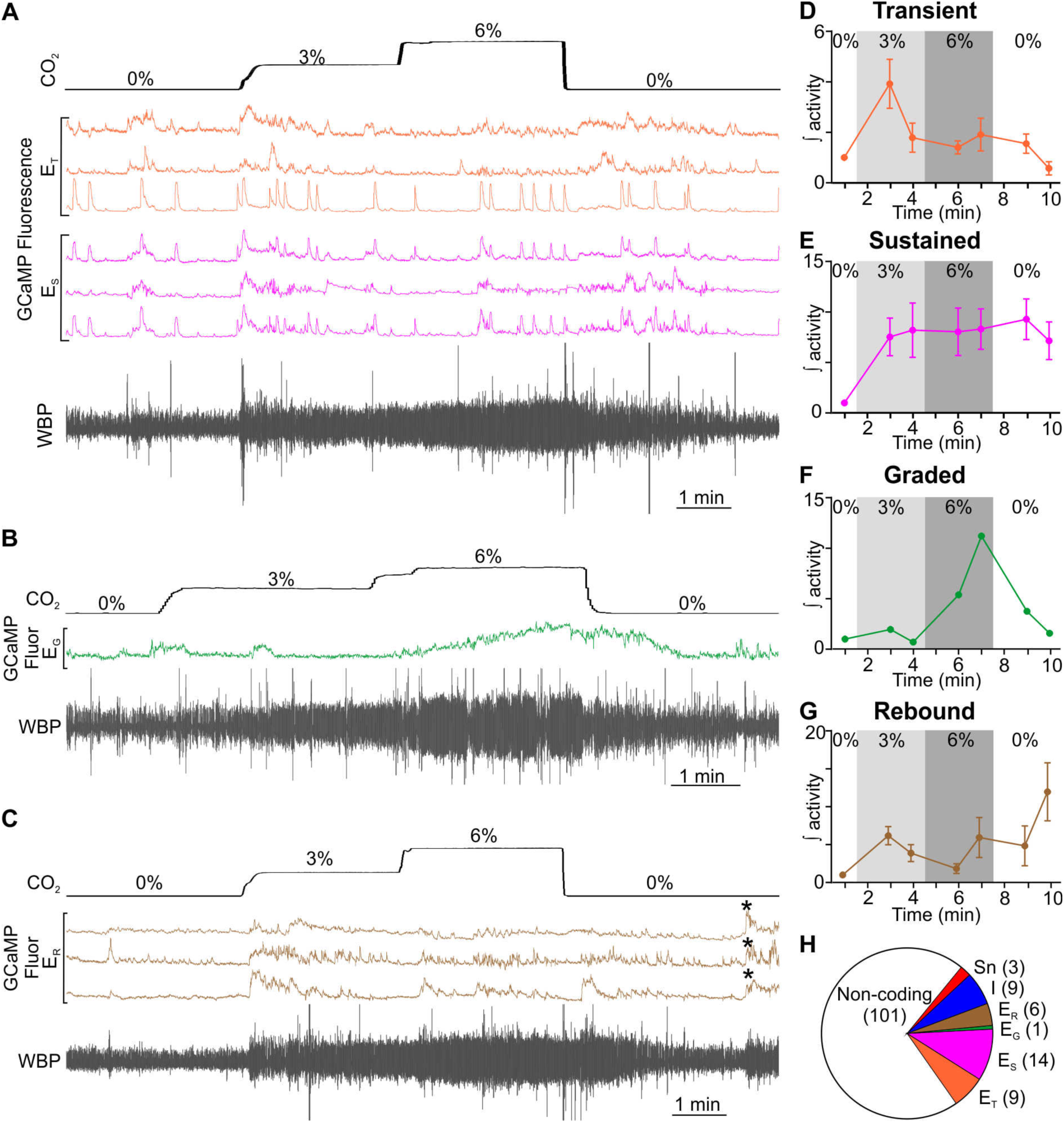
Hypercapnia induced heterogenous RTN neuronal responses. (A-C) WBP recording in response to hypercapnia time-matched with Inscopix recorded GCaMP6 signals. GCaMP6s fluorescence of (A) transient (E_T_; orange) and sustained (E_s_; pink) neurons, (B) graded (E_G_: green) neurons and (C) rebound (E_R_; brown, asterisks show the rebound events) neurons. (D-G) Changes in activity (F/F_0_) of neurons during hypercapnia. (D) Transient, (E) sustained, (F) graded and (G) rebound neurons. (H) Proportion and distribution of recorded RTN neurons into their subtypes.

In summary, for RTN neurons: 101 out of 143 did not encode any aspect of the hypercapnic stimulus; 3/143 encoded sniffing; and the remainder (39/143) had activity patterns that were modulated by CO_2_ (Fig 2H). However these responses were highly heterogeneous, with only one neuron acting like a classical sensory cell and having a response that graded with stimulus strength (E_G_). Given the prior expectation that RTN neurons should exhibit robust changes in firing to CO_2_ ^4-8^, the very low frequency of CO_2_-activated neurons was surprising. This was not due to weak responses to hypercapnia which were very robust in the awake mice (note the raw whole body plethysmography [WBP] traces in Figs 1, 2, and Supplementary Fig 2A). Nor was it due to a technical issue that prevented us from seeing neural activity, as induction of a seizure in anesthetised mice gave clear Ca^2+^ activity in the RTN neurons when seizures occurred and altered respiratory activity (Supplementary Fig 3). We are forced to conclude that the great majority neurons in the RTN are not primarily directly sensing CO_2_. A potential resolution of our conflicting new data with that already in the literature is that we find RTN neurons, unresponsive to CO_2_ in awake animals, are capable of being activated during hypercapnia under anaesthesia (Supplementary Fig 4). The heterogeneous responses are what might be expected if RTN neurons were extracting features related to hypercapnia by integrating the activity of other sensory cells. This is consistent with recent reports of functional heterogeneity in the RTN ^20,21^.

**Figure 3:**
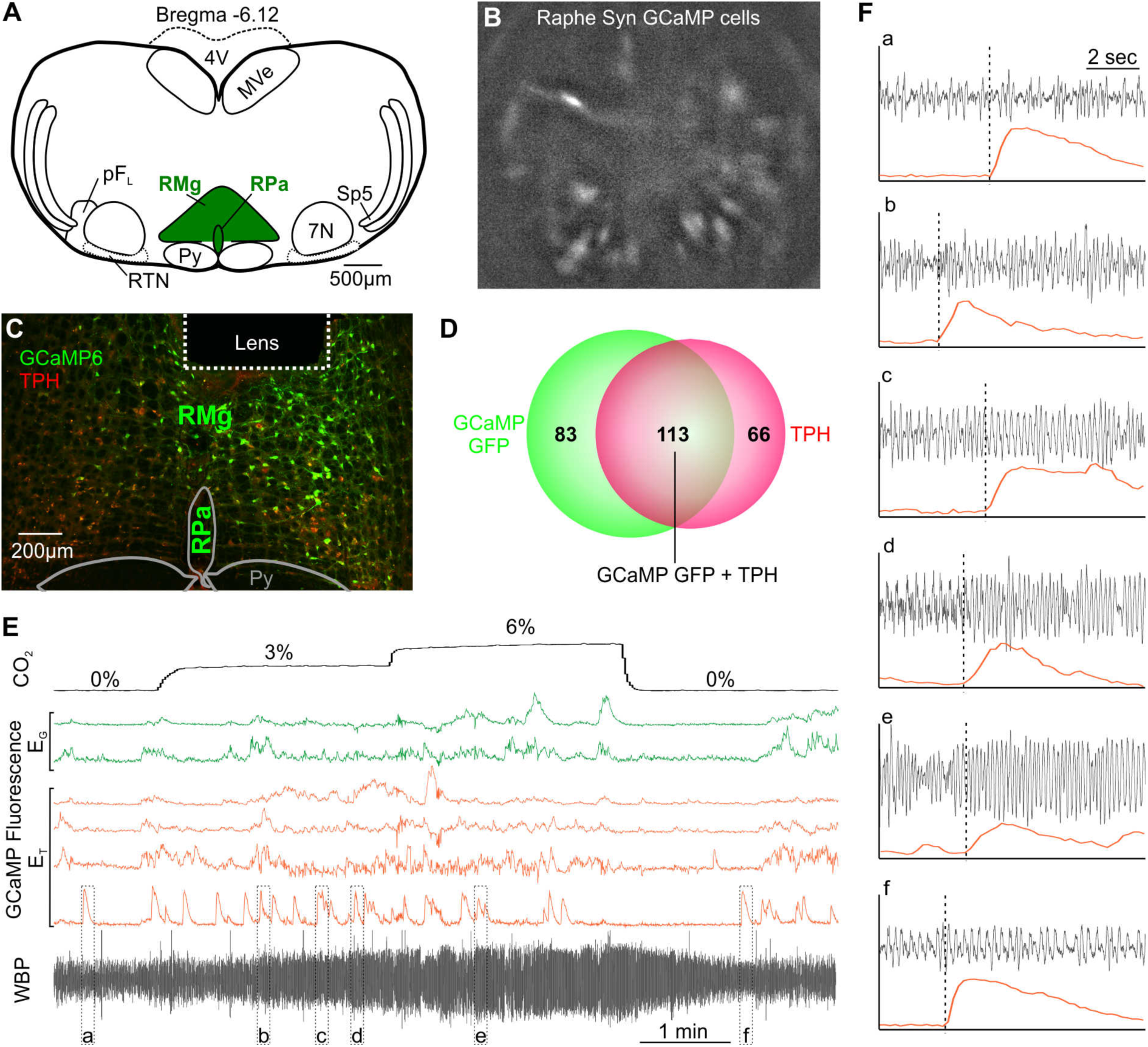
CO_2_-responsive medullary raphe neurons tranduced with synapsin GCaMP6. (A) AAV9-Syn-GCaMP6s injection into the Raphe. (B) GCaMP6s fluorescence signal from GCaMP6s transduced raphe neurons in freely behaving mice. (C) Micrograph of lens placement and viral transduction of neurons (green) relative to the Raphe (TPH+ neurons - red). (D) Venn diagram of cell counts of GCaMP6s transduced neurons (green) under the GRIN lens with TPH+ neurons (red) in the raphe. (E) WBP recording in response to hypercapnia time-matched with Inscopix recorded GCaMP6 signals. GCaMP6s fluorescence from transient (E_T_: orange) and graded (E_G_: green) neurons. (F) Expanded recordings from (E), the start of the Ca^2+^ transient is shown by the dotted line. Abbreviations defined in Figure 1.

**Figure 4:**
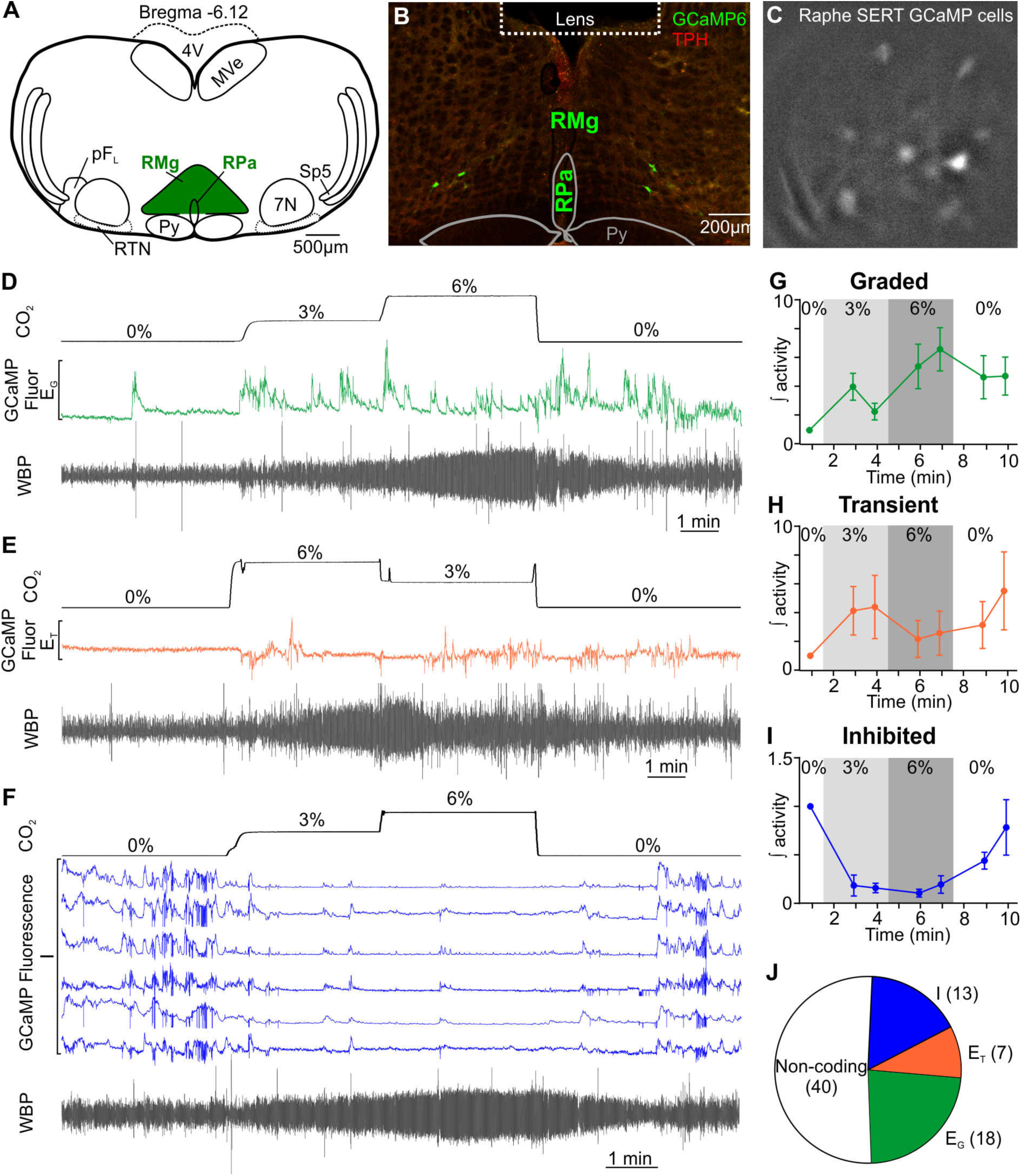
Chemosenstory responses of promoter specific (SERT) raphe neurons to CO_2_ in freely behaving mice. (A) AAV-DJ-SERT-GCaMP6s injection into the Raphe. (B) Micrograph of lens placement and viral transduction of neurons (green) relative to the Raphe (TPH+ neurons - red). (C) GCaMP6s fluorescence signal from transduced raphe neurons in freely behaving mice. (D-F) WBP recording in response to hypercapnia time-matched with Inscopix recorded GCaMP6 signals. GCaMP6s fluorescence of (D) graded (E_G_: green) neurons, (E) transient (E_T_; orange) neurons, and (F) CO_2_-inhibted (I; blue) neurons. (G-I) Changes in activity (F/F_0_) of (G) graded, (H) transient and (I) inhibited neurons during hypercapnia. (J) Proportion and distribution of recorded raphe neurons (both synapsin and SERT specific) into their subtypes. Abbreviations defined in Figure 1.

### Chemosensory responses in the medullary raphe

As the evidence for chemosensory neurons in the rostral medulla is compelling and has been amassed over many years from many independent studies ^3,22-30^, we next examined whether the activity of neurons in the rostral medullary Raphe (i.e. Raphe magnus and pallidus) could be altered by CO_2_ (Figs 3, 4). Once again, we found heterogeneity of responses, but in the rostral medullary Raphe CO_2_-responsive neurons fell into only 3 categories (E_T_, E_G_ and I).

We drove expression of GCaMP6s with a synapsin promoter (n=3; Fig 3A-C). Assessing the ability of this construct to transduce serotonergic neurons in this nucleus, we found that 58% of the transduced neurons in the optical pathway of the microscope were TPH+ (tryptophan hydroxylase; a marker of serotonergic neurons) (Fig 3D). As there are also GABAergic neurons ^31-33^, and non-serotonergic NK1R neurons in the Raphe ^33,34^, we also specifically targetted the serotonergic neurons by driving GCaMP6 expression with a SERT (slc27a4) promoter (n=4, Fig 4). Post-hoc immunocytochemistry showed that all neurons in the optical pathway of the microscope (Fig 4B) which expressed GCaMP6 co-localised with TPH. We recorded 47 synapsin GCaMP6 neurons in 3 mice and 31 SERT GCaMP6 neurons in 4 mice.

7/78 of the responding neurons were transiently activated by 3% inspired CO_2_ and were thus of the E_T_ subtype (Fig 3E and Supplementary Movie 5, organge traces, synapsin; Fig 4E and Supplementary Movie 6, SERT; Fig 4H). 18/78 of the responding neurons were of the E_G_ subtype and showed a graded response to 3 and 6% inspired CO_2_ (Fig 3E and Supplementary Movie 5, green traces, synapsin; Fig 4D and Supplementary Movie 7, SERT; Fig 4G). The Ca^2+^ responses during hypercapnia also exhibited heterogeneity. Some neurons exhibited periodic Ca^2+^ transients, suggestive of burst firing (Fig 3E, F; Fig 4D), whereas others exhibited more sustained elevations of Ca^2+^ suggestive of high frequency unpatterned firing. Surprisingly, 13/78 responding neurons (including serotonergic neurons) were inhibited during hypercapnia (Fig 4F, I and Supplementary Movie 8). A far greater proportion of neurons (38/78, 7 mice; Fig 4J) responded to CO_2_ than in the RTN (χ^2^=10.22, p=0.00139).

Overall, the medullary Raphe appears to possess many more neurons than the RTN that encode the hypercapnic stimulus in a proportional fashion. Thus it seems that the Raphe has more characteristics of a primary chemosensory nucleus than the RTN.

### Chemosensory responses in glial cells

As there is evidence for glial cells acting as chemosensory cells, we transduced mice with an AAV-GFAP-GCaMP6s to drive expression of GCaMP in astrocytes in both the RTN and Raphe (Fig 5A-D). The glial cell fluorescence was more diffuse than for neurons (Fig 5B, D), this is consistent with astrocytes having a large arborisation of tiny processes. Responding astrocytes were often seen adjacent to blood vessels in both areas (Fig 5B, D).

**Figure 5:**
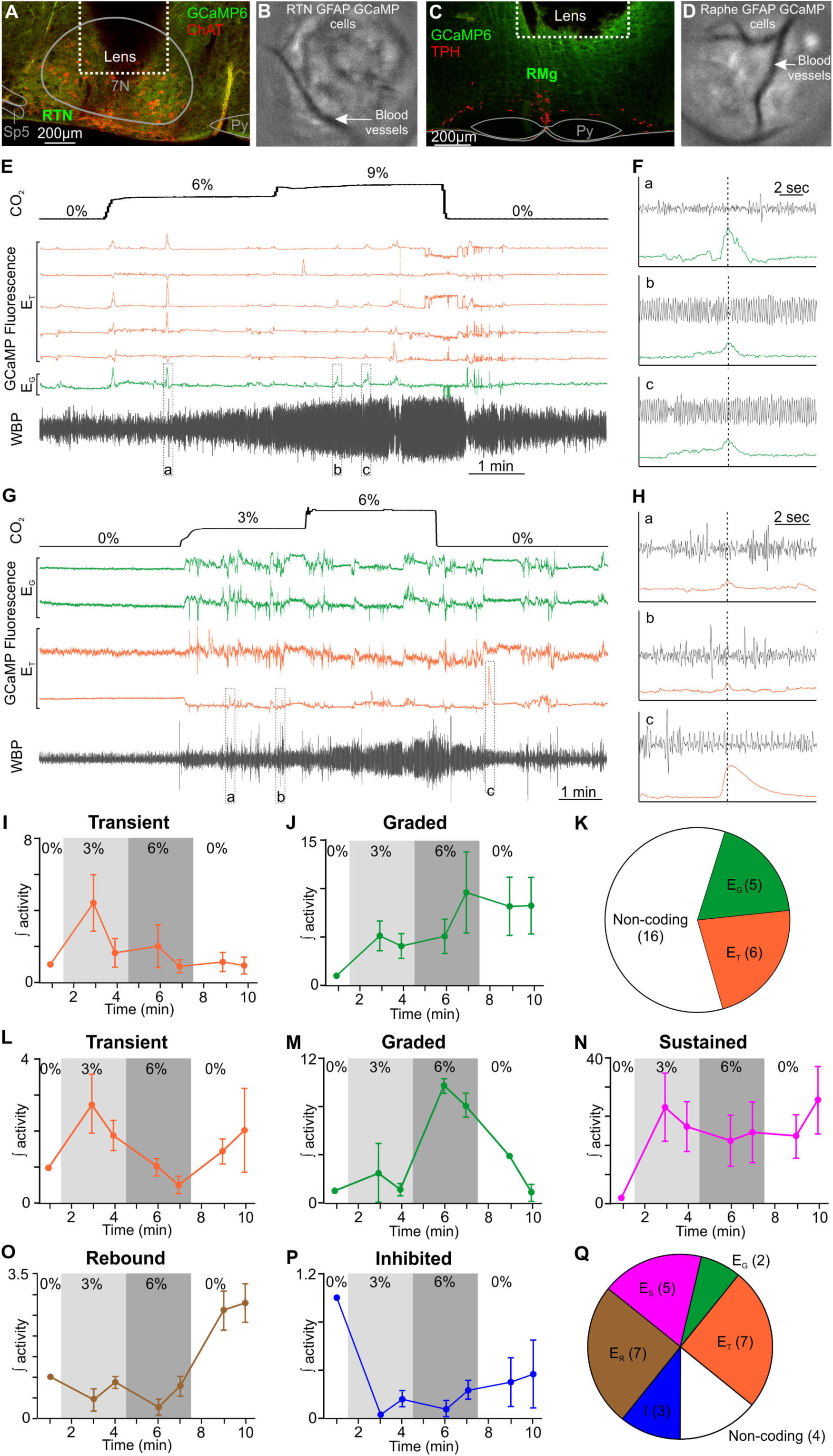
RTN and raphe glia elicit various responses to elevated CO_2_. (A) Micrograph of lens placement and viral transduction of RTN glia (green) relative to the facial nucleus (ChAT+ neurons - red). (B) GCaMP6s fluorescence signal from transduced RTN glial cells in freely behaving mice. (C) Micrograph of lens placement and viral transduction of Raphe glia (green) relative to the Raphe (TPH+ neurons - red). (D) GCaMP6s fluorescence signal from transduced Raphe glial cells in freely behaving mice. (E) WBP recording in response to hypercapnia time-matched with Inscopix recorded GCaMP6 signals. GCaMP6s fluorescence of transient (E_T_; orange) and graded (E_G_: green) RTN glia. (F) Expanded trances from (E), the start of the glial Ca^2+^ transient is shown by the dotted line. (G) WBP recording in response to hypercapnia time-matched with Inscopix recorded GCaMP signals. GCaMP6s fluorescence of transient (E_T_; orange) and graded (E_G_: green) Raphe glia. (H) Expanded trances from (G), the start of the glial Ca^2+^ transient is shown by the dotted line. (I-J) Changes in activity (F/F_0_) of RTN (I) transient and (J) graded glia during hypercapnia. (K) Proportion and distribution of recorded RTN glia. (L-P) Changes in activity (F/F_0_) of Raphe (L) transient, (M) graded, (N) sustained, (O) rebound, and (P) inhibited glia during hypercapnia. (Q) Proportion and distribution of recorded Raphe glia into their subtypes. Abbreviations defined in Figure 1.

In the RTN 11 out of 27 glial cells exhibited a response to hypercapnia (Fig 5E, F, I-K). In 6/27, the response was transient at the beginning of the 3% CO_2_ stimulus (Fig 5E and Supplementary Movie 9, E_T_, orange traces), the remaining 5/27 gave graded responses (Fig 5E and Supplementary Movie 9, E_G_, green trace). These responses in RTN glia are consistent with previous reports of their contribution to the chemosensory control of breathing ^29^. The Ca^2+^ transients in RTN glial cells did not obviously correlate with any specific feature of the plethysmography trace (Fig 5F).

In the Raphe 24 out of 28 glial cells responded to hypercapnia (Fig 5G, H, L-Q). The Raphe glial responses were also heterogenous (Fig 5G, L-P). 7/28 of the responding glial cells gave transient responses (Fig 5G, L and Supplementary Movie 10, E_T_, orange traces), 2/28 were graded (Fig 5G, M and Supplementary Movie 10, E_G_, green traces), 5/28 were sustained (Fig 5N, E_S_, pink trace), 7/28 gave rebound response (Fig 5O, E_R_, brown trace), and 3/28 were inhibited (Fig 5P, I, blue trace). The Ca^2+^ transients in Raphe glia did not consistently correspond to events on the plethysmography trace (Fig 5H). For both the RTN and Raphe transduced mice, robust adaptive ventilatory responses to hypercapnia were seen (Supplementary Fig 2C, D). Thus although glial cells in both the RTN and Raphe, respond to hypercapnia, the proportion of responding cells is higher in the Raphe (χ^2^=16.96, p=0.00004). Interestingly, the proportion of responding glial cells in both nuclei (RTN 41%, Raphe 86%) is higher than the proportion of responding neurons (RTN 26%, Raphe, 48%), demonstrating the potential importance of glia in mediating chemosensory responses to CO_2_.

### Activity of neurons in the pF_L_

Neurons of the pF_L_, lateral and adjacent to the RTN, have been proposed as the expiratory oscillator ^13-15^. Under resting conditions eupneic breathing, with the exception of during REM sleep ^35^, does not involve active expiration, so the neurons of the pF_L_ are largely silent. A hypercapnic stimulus causes an increase in respiratory frequency, tidal volume and also the onset of active expiration ^14,36,37^. We therefore transduced neurons of the pF_L_ in 3 mice with the synapsin-GCaMP6f construct (Fig 6A-C) and observed their responses during a hypercapnic stimulus (Fig 6D, E). We observed that the Ca^2+^ activity reflected changes in respiratory frequency with surprising fidelity (compare gray trace (*f*R) with GCaMP traces in Fig 6D). Closer inspection revealed the temporal relationship between Ca^2+^ activity and changes in the plethysmographic traces (Fig 6E, F, H). Ca^2+^ activity in the pF_L_ neurons on average preceded an increase in tidal volume and respiratory frequency by ∼0.4 s, though some neurons showed Ca^2+^ activity after changes in tidal volume and respiratory frequency had occurred. This time separation was independent of inspired CO_2_. Activity in the pF_L_ neurons always preceded active expiration. At 6% inspired CO_2_ this time difference was ∼0.5 s, while at 9% inspired CO_2_ the time interval shortened to ∼0.3 s (Fig 6I, J). Overall 20/31 neurons in the pF_L_ exhibited responses that temporally preceded active expiration (Fig 6K). Post-hoc immunocytochemistry showed that a small population of neurons which expressed GCaMP6 co-localised with ChAT (11%, Fig 6G) and were located in the optical pathway of the microscope (Fig 6B).

**Figure 6:**
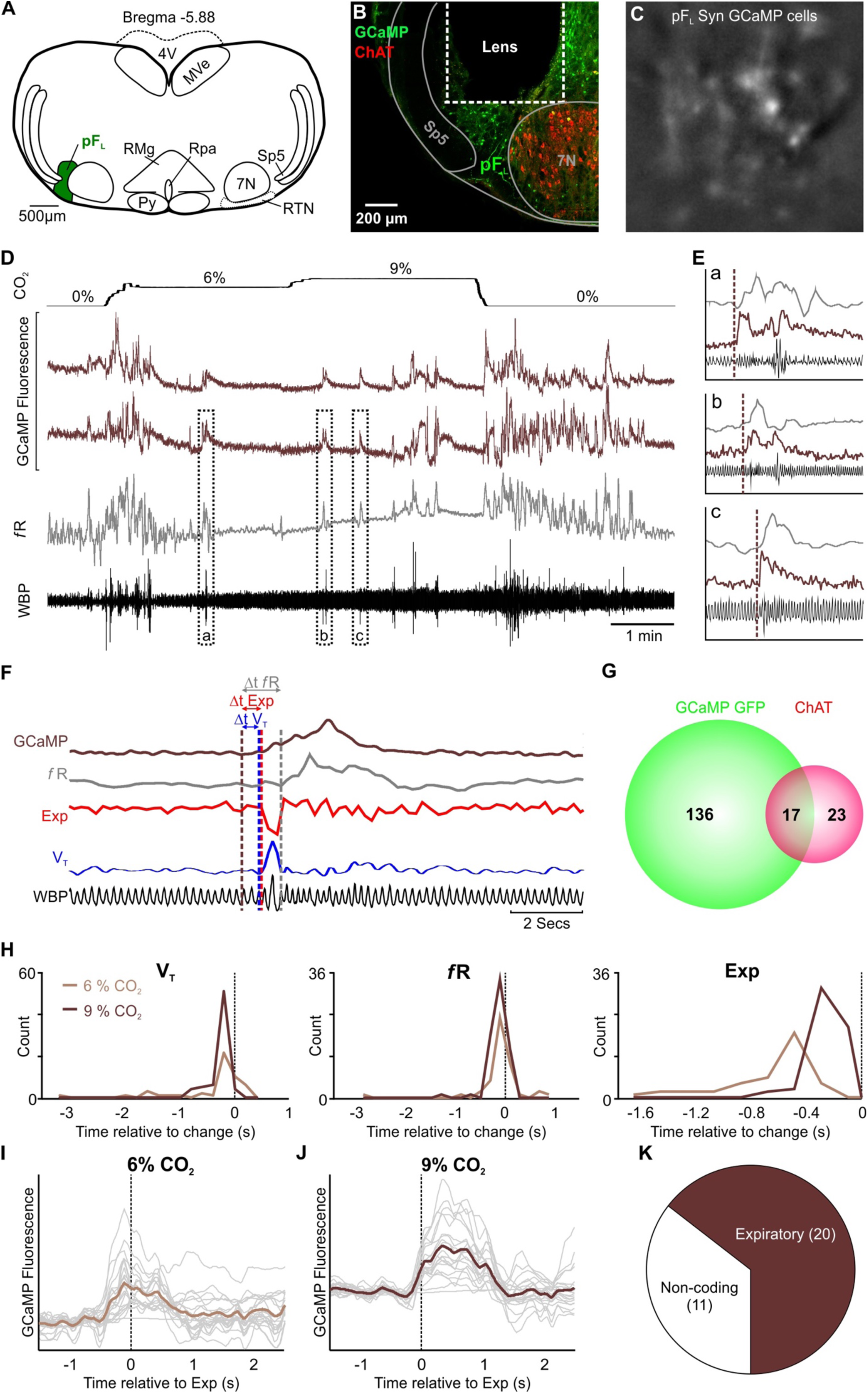
pF_L_ neurons drive active expiration. (A) AAV9-Syn-GCaMP6f injection into the pF_L_. (B) Micrograph of lens placement and viral transduction of pF_L_ neurons (green) relative to the facial nucleus (ChAT+ neurons - red). (C) GCaMP6f fluorescence signal from transduced pF_L_ neurons in freely behaving mice. (D) WBP recording in response to hypercapnia time-matched with Inscopix recorded GCaMP signals. GCaMP6f fluorescence of transient expiratory (Exp; dark brown) pF_L_ neurons. (E) Expanded traces from (D), the start of the Ca^2+^ transient is shown by the brown dotted line. (F) Measurements of GCaMP6 fluorescence (dark brown) relative to tidal volume (V_T_, blue), respiratory frequency (*f*R, grey), and expiration (Exp, red). Verticle dotted lines represent the start of the changes on their respective channels. (G) Venn diagram of cell counts of GCaMP6f transduced (green) and ChAT+ (red) neurons under the GRIN lens. (H) Frequency histograms of timing of changes in GCaMP fluorescence of pF_L_ neurons relative to tidal volume (V_T_, left), respiratory frequency (*f*R, centre), and expiration (Exp, right). (I-J) Spike triggered average (brown line) of all calcium events (grey lines) temporally correlated to the beginning of expiratory activity (dotted verticle line) at (I) 6% CO_2_ and (J) 9% CO_2_. (K) Proportion of recorded pF_L_ neurons that showed the expiratory activity. Abbreviations defined in Figure 1.

**Figure 7:**
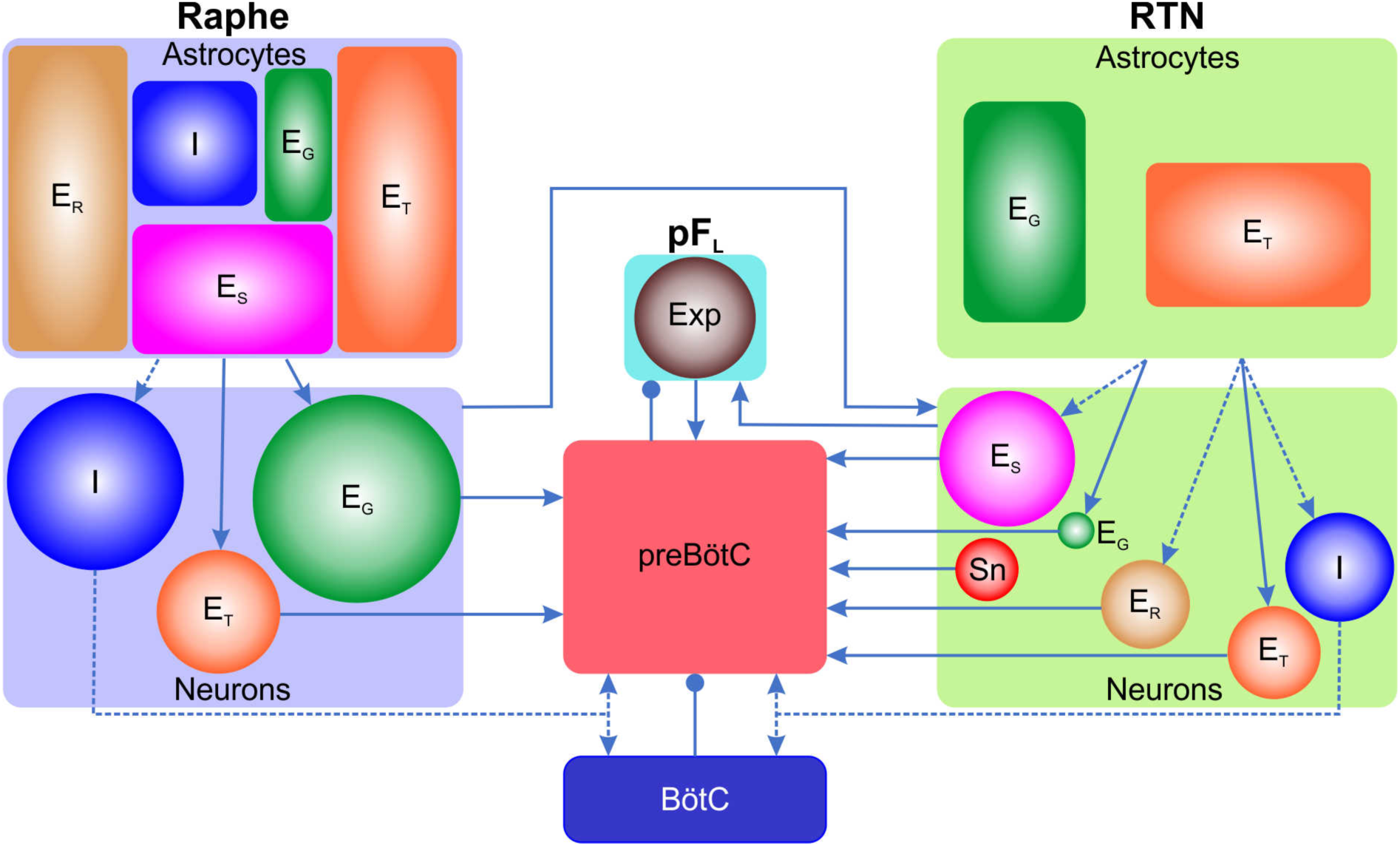
Summary model for the contribution of the RTN, Raphe and pF_L_ neurons and glia to respiratory chemosensitivity and active expiration. Neurons are shown as circles. Astrocytes are shown as rectangles. The areas of each rectangle and circle shows the proportion of each type of response observed. Note that for both astrocytes and neurons a far higher proportion of cells respond in the Raphe compared to the RTN. Cells excited by CO_2_ (E_T_, E_S_, E_R_, E_G_) are hypothesized to project directly to the preBötC. Cells inhibited by CO_2_ are hypothesized to project to (and excite) the BötC. CO_2_ would thereby remove tonic excitation from BötC neurons (which inhibits the preBötC) and thus disinhibit neurons in the preBötC. Alternatively CO_2_-inhibited neurons could project directly to the preBötC and excite inhibitory interneurons therein. During hypercapnia, this excitation would weaken, reducing levels of inhibition within the preBötC network and allowing coordination of stronger inspiratory bursts. Glia are hypothesized to excite their equivalent neuronal counterparts and thus comprise a key part of the chemosensory network. The Raphe project to the RTN and provides chemosensory drive to this nucleus ^44^. The RTN excites neurons of the pF_L_ to drive active expiration ^64^. The pF_L_ is coupled to the preBötC via excitation from pF_L_ to preBötC and reciprocal inhibitory connections from preBötC to pF_L_ ^15^. Abbreviations: E_G_ -excited with a graded response to CO_2_; E_T_ -excited with a transient respone to CO_2_; E_S_ -excited with a sustained constant response to CO_2_; E_R_ -excited on rebound after hypercapnia; I -inhibited by CO_2_; Sn -sniff-coding; Exp -expiratory active neurons. Arrow head -excitation; round head -inhibition.

## Discussion

### Chemosensory mechanisms in adult mice

Evidence for the involvement of the RTN, and in particular the Phox2B+ neurons, in CO_2_ chemoreception comes mainly from neonatal or young juvenile animals ^6-8^. This prior evidence depends heavily on recordings from anaesthetized animals. There is also evidence that by about 3 months chemosensory control adapts and has much less dependence on the Phox2B+ neurons of the RTN ^7^. The highly heterogeneous responses of RTN neurons to a hypercapnic stimulus was therefore very surprising. Although some neurons indicated the presence of hypercapnia, only 1/143 neurons was of the type that might have been expected from the prior literature: i.e. exhibiting a graded response to CO_2_, that encoded the magnitude of the hyperapnic stimulus. Unexpectedly there were some neurons that were inhibited by hypercapnia. Other neurons appeared to encode features of hypercapnia (such as its start and end) rather than its magnitude. This heterogeneity is reminiscent of other sensory systems, e.g. the auditory, where neurons in secondary nuclei (e.g. the cochlear nucleus) show a range of responses that abstract features of the primary auditory stimulus. We suggest that the available evidence suggests that the majority of neurons that we recorded from the RTN were not directly sensing CO_2_, but rather integrating responses from other directly chemosensitive cells, such as the Raphe ^38^ and peripheral chemoreceptors ^39^.

Reconciliation with the prior literature is possible through our observation that anaesthesia enhances CO_2_ responses in RTN neurons (despite weakening the overall hypercapnic response, Supplementary Fig 4). The effect of anesthesia in reconfiguring the operation of brainstem neuronal circuits may therefore account for why in adult mice RTN neurons are thought to play a central role in chemoreception. Our recordings have been performed in awake adult mice, and it remains probable that RTN neurons are indeed central to the chemosensitivity of neonatal mice ^7,40^. Recently the pH sensitive channel GPR4 has been claimed to endow RTN neurons with pH sensitivity ^8^. The extent to which GPR4 in RTN neurons contributes to respiratory chemosensitivity is the subject of conflicting evidence ^41^. Global knockout of GPR4 severely attenuates the adaptive response to hypercapnia ^8^, but this channel is widely expressed centrally and peripherally including in the carotid body chemosensory cells and the medullary raphe ^8,41^.

Several lines of evidence have suggested the importance of medullary Raphe serotonergic neurons in mediating respiratory chemosensitivity: the proximity of raphe neuron processes to blood vessels ^42^; the correspondence of the location of CO_2_-dependent ATP release at the medullary surface and the raphe neurons ^11,28^; suppression of the medullary raphe by isofluorane ^12,43^ which matches the reduction in the overall hypercapnic ventillatory response; and most compellingly the observation that inhibition of their activity via inhibitory DREADD receptors removes ∼40% of the total adaptive ventilatory response in awake mice ^12^. Our *in vivo* recordings from freely moving mice suggest that in the rostral chemosensory area, it is the Raphe neurons that play the major role in mediating chemosensitivity.

Raphe neurons are well placed to have far-reaching effects on breathing. Of the 2 known serotonergic subtypes, NK1R negative serotonergic neurons project to areas responsible for CO_2_ integration ^44^, whereas NK1R positive serotonergic neurons project to motor nuclei responsible for airway patency and co-ordination, and diaphragmatic movements ^34^. Interestingly both subtypes project to the preBötC ^34,44^. Whilst neither sertonergic group project to pF_L_ ^34^, they do innervate areas that influence expiration, namely the C1 ^45^, and NTS ^46^. Furthermore, the GABAergic neurons of the Raphe have further reaching effects than just inhibition of their sertonergic neighbours ^47^, as GAD positive neurons in the Raphe magnus project to the phrenic motor nucleus ^48^, and the rostral ventrolateral medulla ^49^. That medullary Raphe neurons express GPR4, suggests they are intrinsically pH-sensitive. Given their proximity to sites of CO_2_-dependent ATP release ^28,30^, they may also be a site for convergence of direct CO_2_ sensitivity mediated via Cx26 ^50^.

The presence of CO_2_-inhibited neurons in both the RTN and serotonergic subtypes of the Raphe was a further unexpected finding. If these neurons were to tonically excite neurons that normally inhibit the preBötzinger complex, a process of disinhibition during hypercapnia could result in greater excitation of preBötzinger complex neurons. RTN neurons are known to project to: i) the Bötzinger complex ^51^, which contains inhibitory neurons ^52,53^; and ii) inhibitory cells of the preBötC ^54^, which regulate the synchronisation of excitatory bursting and the emergence of the inspiratory rhythm ^55^. The CO_2_-dependent disinhibition of the inhibitory interneurons in the preBötC could contribute to the emergence of more powerful inspiratory bursts and hence an increase in tidal volume.

### Non-chemosensory mechanisms in the RTN control sniffing

Sniffing is a mechanism that allows for olfactory sampling of the air to discern location of smell and to test for irritants before air is taken into the lung. Sniffing is obligatory for odour perception ^56^, and thus activates the piriform cortex ^57,58^. However, sniffing also causes neuronal activity in the absence of odorants ^58^. It activates the hippocampus ^59,60^, and the cerebellum ^61^, respectively likely to prime odour-related memory recall, and co-ordination of movement towards or away from specific odours. Whilst the activation of these brain regions during sniffing have been well documented, the pattern generating microcircuit for sniffing has only recently been identified. Sniffing requires the activation of 2 sets of muscles i) the nasal muscles to open the airway and decrease resistance, allowing more rapid movement of air through the nose, and ii) the respiratory (inspiratory and expiratory) muscles, to draw air in and out of the nasal cavity. The RTN is located adjacent to, and innervates, the facial motor nucleus ^62,63^. These RTN projections to the facial are directly involved in sniffing ^63^ and control of nasal direction during the odour response ^62^. In conjunction the RTN provides significant drive to the preBötC ^51^, and the pF_L_ ^64^, both of which are integral parts of the sniffing microcircuit ^63,65^. Therefore it is not surprising that the sniff pathway originates in, or passes through, the RTN.

### Role of glial cells

Glial cells have recently been implicated in chemosensory responses. Within the RTN, astrocytes can control breathing via pH-dependent ATP release ^37^. Glial cells in the ventral medulla also express Cx26 ^30^, which is intrinsically sensitive to CO_2_ ^66,67^ and contributes to the adaptive ventilatory response to CO_2_ ^30^. Our recordings from glial cells in both the RTN and Raphe support the hypothesis that these cells have an important role in mediating chemosensory reflexes. In both areas there was a higher proportion of glial cells that responded to hypercapnia than there was for neurons. Strangely, the complexity of glial responses in the Raphe was greater than that of the responses seen in neurons. The opposite was true in the RTN, where the glial responses were much simpler than those exhibited by neurons.

### Active expiration

The pF_L_ is thought to be a subsidiary conditional oscillator for expiration that is normally suppressed at rest ^15^. That all pF_L_ neurons showed the same discharge pattern, supports the hypothesis that the pF_L_ is comprised of a single homogenous group of neurons acting with a single purpose ^20^. Even though the nucleus appears to integrate information from many nuclei -RTN ^64^, C1, ^45^, NTS, rostral pedunculopontine Tegmental ^46^ and the preBötC ^15^ -the pF_L_ appears to be a simple on-off switch, rather than forming complex outputs as we describe in the RTN. Interestingly not all transduced neurons expressed expiratory activity. These remaining neurons are either not recruited during hypercapnia, or instead may either be: i) involved in other physiological reflexes that induces active expiration, e.g. hypoxia ^14,36,45^; or ii) facial neurons, a small population of which expressed GCaMP6 (Fig 6G).

Surprisingly, we found that pF_L_ neurons were only transiently active during large expiratory efforts, (such as sniffing, sighing, or other forms of brief deep breathing), in contrast to the sustained active expiration previously reported ^14,68^. This may be due to our use of brief periods of graded CO_2_ in conscious animals, rather than sustained CO_2_ (∼1 hour) ^68^ or use of the vagotomised anaesthetised preparation in prior studies ^14^. Furthermore, in contrast to our expectation that pF_L_ neurons would be phasically active between inspirations, pF_L_ neurons exhibited elevated intracellular calcium beginning before, and spanning, the expiratory period. Therefore, the phasic response seen in the abdominal muscles almost certainly occurs through inhibition of the motor output mediated by the preBötC during inspiration, providing an explanation as to why the pF_L_ is unable to drive phasic expiratory activity in the absence of rhythmic output from the preBötC ^15^.

### Concluding remarks

Endoscopic imaging of the activity of the brainstem neurons and glia involved in the control of breathing provides a new way to investigate the neural circuitry for the chemosensory control of breathing in awake adult mice. Some components, such as the expiratory activity of the pF_L_ or sniff activity in the RTN, are easy to interpret, whilst others are more complicated. The neuronal responses to CO_2_ were highly heterogeneous in both the RTN and Raphe, however a higher proportion of neurons in the Raphe had the hallmarks of what would be expected of classical chemosensory neurons. This suggests that the Raphe is the primary site of chemoreception and, in the adult, the RTN is a secondary relay nucleus that abstracts features of the chemosensory stimulus such as its start and end. Somewhat surprisingly, a higher proportion of glia responded to hypercapnia than neurons suggesting these cells are important components of the chemosensory mechanisms. Whilst collectively glia displayed the same categories of response to hypercapnia as neurons, the responses of glia did not completely mirror those of neurons in the same nucleus. In the RTN where glia showed fewer response types, it is most likely that glia whose activity matched neuronal response (i.e., E_T_ and E_G_) are either driving the neurons through release of gliotransmitter, or are activated by ionic changes when RTN neurons depolarise. However in the Raphe, glial responses were more complex than their neuronal neighbours, ruling out the possibility glia are simply following neuronal activity or vice versa. This opens up the possibility that glia and neurons work synergistically to process and integrate the salient signals in primary chemosensory nuclei and prompts the question, what is the need for such diversity?

## Materials and Methods

Experiments were performed in accordance with the European Commission Directive 2010/63/EU (European Convention for the Protection of Vertebrate Animals used for Experimental and Other Scientific Purposes) and the United Kingdom Home Office (Scientific Procedures) Act (1986) with project approval from the University of Warwick’s AWERB.

### AAV-DJ: pAAV-SERT-GCaMP6s Vector particle production and purification

The CMV promoter region was excised from pAAV-GFP (Cellbiolabs) by restriction digest with MluI and ClaI, blunting with Klenow Fragment (TheroScientific) and re-ligation with T4 DNA ligase (ThermoScientific) to generate pAAV-GFP without promoter. A SERT promoter clone was purchased from Genecopoiea (Product ID: MPRM41232-PG02, Symbol: Slc6a4, Species: Mouse, Target Gene Accession: NM_010484, Alias: 5-HTT, AI323329, Htt, SertA). The promoter region was cut out with EcoRI and HndIII and introduced by T4 DNA ligation into the previously modified pAAV-GFP to generate pAAV-SERT. Subsequently, a PCR performed using PrimeStar GLX Polymerase (Takara Clontech) with pAAV.Syn.GCaMP6s.WPRE.SV40 (Addgene Plasmid#100843) as a template (forward primer 5’- TTGACTGCCTAAGCTTgccaccatgcatcatcatcatcatcatg and reverse primer 5’- GATCTCTCGAGCAGCGCTtcacttcgctgtcatcatttgtacaaact) to amplify the GCaMP6s fragment. pAAV-SERT was digested with HindIII and AfeI and the PCR product was inserted by InFusion Cloning (Takara Clontech) to generate pAAV-SERT His-GCaMP6s. Accuracy of all cloning steps was verified by PCR, restriction digests and DNA sequencing analysis.

AAV-DJ pseudotyped vectors were generated by co-transfection of HEK293-AAV cells (Cellbiolabs) with the AAV transfer plasmid pAAV-SERT His-GCaMP6s and the AAV packaging plasmid pAAV-DJ and pHelper (both Cellbiolabs). HEK293-AAV cells (Cellbiolabs) were cultivated in Dulbecco’s modified Eagle’s medium (DMEM, High Glucose, Glutamax) supplemented with 10% (v/v) heat-inactivated fetal calf serum, 0.1 mM MEM Non-Essential Amino Acids (NEAA), 100 U/ml penicillin and 100 μg/ml streptomycin. Tissue culture reagents were obtained from Life Technologies. Briefly, 1×10^7^ HEK293-AAV cells were seeded one day before transfection on 15-cm culture dishes and transfected with 7.5 µg pAAV-DJ, 10 µg pHelper and 6.5 µg pAAV plasmid per plate complexed with Max-polyethylenimine (PEI, Polysciences) at a PEI:DNA ratio (w/w) of 3:1 {Fukumoto, 2010 #5700}. After 72 h cells were harvested and resuspended in 5 ml lysis buffer (50 mM Tris base, 150 mM NaCl, 5 mM MgCl_2_, pH 8.5). After three freeze-thaw cycles, benzonase (Merk; final concentration 50 U/ml) was added and the lysates were incubated for 1 h at 37°C. Cell debris was pelleted and vector containing lysates were purified using iodixanol step gradients. Finally, iodixanol was removed by ultrafiltration using Amicon Ultra Cartridges (50 mwco) and three washes with DPBS. The genomic titers of DNase-resistant recombinant AAV particles were determined after alkaline treatment of virus particles and subsequent neutralization by qPCR using the qPCRBIO SY Green Mix Hi-Rox (Nippon Genetics Europe GmbH) and an ABI PRISM 7900HT cycler (Applied Biosystems). Vectors were quantified using primers specific for the GCaMP6s sequence (5’-CACAGAAGCAGAGCTGCAG and 5’-actggggaggggtcacag). Real-time PCR was performed in a total volume of 10 μl with 0.3 μM for each primer. The corresponding pAAV transfer plasmid was used as a copy number standard. A standard curve for quantification was generated by serial dilutions of the respective plasmid DNA. The cycling conditions were as follows: 50°C for 2 min, 95°C for 10 min, followed by 35 cycles of 95°C for 15 sec and 60°C for 60 sec. Calculations were done using the SDS 2.4 software (Applied Biosystems).

### Viral handling

Raphe and RTN neurons - AAV-9: pGP-AAV-syn-GCaMP6s-WPRE.4.641 at a titre of 1×10^13^ GC·ml- ^1^ (Addgene, Watertown, MA, USA);

Raphe: AAV-DJ: pAAV-SERT-GCaMP6s at a titre of 1.8×10^13^GC·ml^-1^ (University hospital Hamburg-Eppendorf, Hamburg, Germany).

pF_L_ - AAV-9: pGP-AAV-syn-GCaMP6f-WPRE.24.693 at a titre of 1×10^13^ GC·ml^-1^ (Addgene, Watertown, MA, USA).

Raphe and RTN glia AAV-5: pZac2.1 gfaABC1D-cyto-GCaMP6f at a titre of 1×10^13^ GC·ml^-1^ (Addgene, Watertown, MA, USA).

Viruses were aliquoted and stored at −80°C. On the day of injection, viruses were removed and held at 4°C, loaded into graduated glass pipettes (Drummond Scientific Company, Broomall, PA, USA), and placed into an electrode holder for pressure injection. The AAV-syn-GCaMP6s and AAV-syn-GCaMP6f vectors use the synapsin promoter, and therefore transduced neurons showing higher tropism for the AAV 2/9 subtype, and to a much lesser extent neurons that show low tropism for AAV 2/9 within the injection site, e.g. facial motoneurons, they do not transduce non-neuronal cells. Whereas vector AAV SERT-GCaMP6s and AAV GFAP-GCaMP6f were specific for serotonergic neurons and astrocytes, respectively.

### Viral transfection of RTN, Raphe neurons and astrocytes, and pF_L_ neurons

Adult male C57BL/6 mice (20-30 g) were anesthetized with isofluorane (4%; Piramal Healthcare Ltd, Mumbai, India) in pure oxygen (4 L·min^-1^). Adequate anaesthesia was maintained with 0.5-2% Isofluorane in pure oxygen (1 L·min^-1^) throughout the surgery. Mice received a presurgical subcutaneous injection of atropine (120 µg·kg^-1^; Westward Pharmaceutical Co., Eatontown, NJ, USA) and meloxicam (2 mg·kg^-1^; Norbrook Inc., Lenexa, KS, USA). Mice were placed in a prone position into a digital stereotaxic apparatus (Kopf Instruments, Tujunga, CA, USA) on a heating pad (TCAT 2-LV: Physitemp, Clifton, NJ, USA) and body temperature was maintained at a minimum of 33°C via a thermocouple. The head was levelled, at bregma and 2 mm caudal to bregma, and graduated glass pipettes containing the virus were placed stereotaxically into either the RTN, rostral medullary raphe or pF_L_ (Fig 1A, 3A, 4A and 6A). The RTN was defined as the area ventral to the caudal half of the facial nucleus, bound medially and laterally by the edges of the facial nucleus (coordinates with a 9° injection arm angle: −1.0 mm lateral and −5.6 mm caudal from Bregma, and - 5.5 mm ventral from the surface of the cerebellum; Fig 1A). The Raphe defined as the medial, TPH containing regions level with the caudal face of the facial nucleus: the Raphe Magnus (RMg) was directly above the pyramidal tracts (Py), and the Raphe Pallidus (RPa) was bound laterally by the Py (coordinates with a 9° injection arm angle: 0 mm lateral and −5.8 mm caudal from Bregma, and - 5.2 mm ventral from the surface of the cerebellum; Fig 3A). The pF_L_ was defined as the neurons bound laterally by the spinothalmic tract and medially by the lateral edge of the facial motor nucleus (coordinates with a 9° injection arm angle: −1.8 mm lateral and −5.55 mm caudal from Bregma, and −4.7 mm ventral from the surface of the cerebellum; Fig 6A). The virus solution was pressure injected (<300 nL) unilaterally. Pipettes were left in place for 3-5 minutes to prevent back flow of the virus solution up the pipette track. Postoperatively, mice received intraperitoneal (IP) injections of buprenorphine (100 µg·kg^-1^; Reckitt Benckiser, Slough, UK). Mice were allowed 2 weeks for recovery and viral expression, with food and water *ad libitum*.

### GRIN lens implantation

Mice expressing GCaMP6 were anesthetized with isofluorane, given pre-surgical drugs, placed into a stereotax, and the head was levelled as described above. To widen the lens path whilst producing the least amount of deformation of tissue, a graduated approach was taken; firstly a glass pipette was inserted down the GRIN lens path to a depth 200 µm above where the lens would terminated and left in place for 3 mins; this procedure was then repeated with a blunted hypodermic needle. The GRIN lens (600 µm diameter, 7.3 mm length; Inscopix, Palo Alto, CA, USA) was then slowly inserted at a rate of 100µm·min^-1^ to a depth ∼1300 µm above the target site, then lowered at a rate of 50µm·min^-1^ to a depth ∼300 µm above the RTN, Raphe or pF_L_ (coordinates with a 9° injection arm angle: RTN – 1.1 mm lateral and −5.75 mm caudal from Bregma, and −5.3 mm ventral from the surface of the cerebellum; Raphe – 0 mm lateral and −5.95 mm caudal from Bregma, and −5.1 mm ventral from the surface of the cerebellum; pF_L_ – 1.7 mm lateral and −5.7 mm caudal from Bregma, and −4.6 mm ventral from the surface of the cerebellum). The lens was then secured in place with SuperBond^(tm)^ (Prestige Dental, Bradford, UK). Postoperatively, mice received buprenorphine, and were allowed 2 weeks for recovery, with food and water *ad libitum*.

### Baseplate installation

Mice expressing GCaMP6 and implanted with GRIN lens were anesthetized with isofluorane, given pre-surgical drugs, and placed into a stereotax as described above. To hold the miniaturized microscope during recordings, a baseplate was positioned over the lens and adjusted until the cells under the GRIN lens were in focus. The baseplate was then secured with superbond^(tm)^, and coated in black dental cement (Vertex Dental, Soesterberg, the Netherlands) to stop interference of the recording from ambient light. Mice were allowed 1 week for recovery, with food and water *ad libitum*.

### Ca^2+^ imaging in freely moving mice

All mice were trained with dummy camera and habituated to plethysmography chamber at least twice before imaging. The miniature microscope with integrated 475 nm LED (Inscopix, Palo Alto, CA, USA) was secured to the baseplate. GCaMP6 fluorescence was visualised through the GRIN lens, using nVista 2 HD acquisition software (Inscopix, Palo Alto, CA, USA). Calcium fluorescence was optimised for each experiment so that the histogram range was ∼150-600, with average recording parameters set at 10-20 frames/sec with the LED power set to 10-20 mW of light and a digital gain of 1.0-4.0. A TTL pulse was used to synchronize the calcium signalling to the plethysmography trace. All images were processed using Inscopix data processing software (Inscopix, Palo Alto, CA, USA). GCaMP6 movies were ran through: preprocessing algorithm (with temporal downsampling), crop, spatial filter algorithm (average 0.005-0.5), motion correction and cell identification through manual regions of interest (ROIs) operation to generate the identified cell sets. Cell sets were imported into *Spike2* software and the activity plot was calculated by the area under curve analysis.

### Movement correction and movement artefacts

The mice were unrestrained and able to move freely during the recordings. Movement artefact evident during most recordings was corrected for via the Inscopix software. High definition videos of mice synchronized to the plethysmograph recordings online in *Spike2* were filmed for visualization of physical movements during analysis. GCaMP6 signals were accepted if the following criteria were met: 1) the features of the cell (e.g. soma, large processes) could be clearly seen; 2) they occurred in the absence of movement of the mouse or were unaffected by mouse movement; 3) fluorescence changed relative to the background; 4) the focal plane had remained constant as shown by other nonfluorescent landmarks (e.g. blood vessels).

### Plethysmography

Mice were placed into a custom-made 0.5 L plethysmography chamber, with an airflow rate of 1 l·min^-1^. The plethysmography chamber was heated to 31^0^C (thermoneutral for C57/BL6 mice). CO_2_ concentrations were sampled via a Hitech Intruments (Luton, UK) GIR250 Dual Sensor Gas analyzer or ML206 gas analyzer (ADinstruments, Sydney, Australia) connected to the inflow immediately before entering the chamber. The analyser had a delay of ∼15-20 sec to read-out the digital output of gas mixture. Pressure transducer signals and CO_2_ measurements were amplified and filtered using the NeuroLog system (Digitimer, Welwyn Garden City, UK) connected to a 1401 interface and acquired on a computer using *Spike2* software (Cambridge Electronic Design, Cambridge, UK). Video data was recorded with *Spike2* software and was synchronised with the breathing trace. Airflow measurements were used to calculate: tidal volume (V_T_: signal trough at the end of expiration subtracted from the peak signal during inspiration, converted to mL following calibration), and respiratory frequency (*f*R: breaths per minute).

### Hypercapnia in freely behaving mice

Instrumented mice were allowed ∼30 mins to acclimate to the plethysmograph. The LED was activated through a TTL pulse synchronised with the *Spike2* recording and 1 or 3 min of baseline recordings were taken (gas mixture: 0% CO_2_ 21% O_2_ 79% N_2_). The mice were then exposed to 3 min epochs of hypercapnic gas mixture at different concentrations of CO_2_ (3, 6 and/or 9%, in 21% O_2_ balanced N_2_), the order of which was randomised. Following exposure to the hypercapnic gas mixtures, CO_2_ levels were reduced back to 0% and calcium signals were recorded for a further 3 minute recovery period.

### Hypercapnia and seizure induction in urethane anaesthetised mice

Instrumented mice were anaesthetised with an IP injection of 1.2-1.5 g·kg^-1^urethane (Sigma-Aldrich, St Louis, MO, USA) and placed into the plethysmograph. For recording responses of RTN neurons to hypercapnia, the LED was activated and 1 minute of baseline recording was taken. The mice were then exposed to 3-minute epochs of hypercapnic gas mixture 3%, 6% and 9% (in 21% O_2_ balanced N_2_), as experiments into the chemosensitivity of RTN neurons in mice often use 6% CO_2_ in freely behaving experiments but 9% CO_2_ in anaesthetised animals. Following exposure to the hypercapnic gas mixtures, CO_2_ levels were reduced back to 0% and calcium signals were recorded for a further 3-minute recovery period.

For neurons recorded in the RTN, after completion of hypercapnic responses and a period of rest, baseline Ca^2+^ activity was recorded for 5 minutes. The mice were then injected with a subthreshold dose of kainic acid to induce behavioural seizures (8 mg·kg^-1^ IP) but sufficient to induce an electrographic seziures. Following injection of kainic acid, calcium activity was recorded every alternate 5 minutes to avoid the fluorophore bleaching and concurrently record the calcium activity for a long enough to evidence the effect of induction of seizure on the activity of RTN neurons (at least for 90 min post kainic acid injection).

### Immunocytochemistry

Mice were humanely killed by pentobarbital overdose (>100 mg·kg^−1^) and transcardially perfused with paraformaldehyde solution (4% PFA; Sigma-Aldrich, St Louis, MO, USA). The head was removed and postfixed in PFA (4°C) for 3 days to preserve the lens tract. The brains were removed and postfixed in PFA (4°C) overnight. Brainstems were serially sectioned at 50-70 μm. Free-floating sections were incubated for 1 hour in a blocking solution (PBS containing 0.1% Triton X-100 and 5% BSA). Primary antibodies (goat anti-cholineacetyl transferase [ChAT; 1:100; Millipore, Burlington, MA, USA], or rabbit anti-tryptophan hydroxylase [TPH; 1:500; Sigma-Aldrich, St Louis, MO, USA] antibody) were added and tissue was incubated overnight at room temperature. Slices were washed in PBS (6 × 5 mins) and then optionally incubated for 1 hour in a blocking solution before the addition of the secondary antibodies: either donkey anti-rabbit Alexa Fluor 680 (1:250; Jackson Laboratory, Bar Harbor, ME, USA), or donkey anti-goat Alexa Fluor 568 (1:250; Jackson Laboratory, Bar Harbor, ME, USA) antibody, tissue was then incubated for 2-4 hours at room temperature. Tissue was washed in PBS (6 × 5 min). Slices were mounted, coverslipped and examined using a Zeiss 880 confocal microscope with ZEN aquisition software (Zeiss, Oberkochen, Germany).

## Author contributions

AB performed experiments and analyzed data; JvdW assisted with surgeries and measurements; RR performed immunocytochemistry; IB created the the SERT viral contruct, RH designed and oversaw the surgical aspects of the study, performed experiments and analyzed data; ND conceived and designed the study; AB, RH and ND reviewed the data and contributed to writing of the manuscript.

## Data and Materials availability

Raw data available on request.

---

**Supplementary Movie 1: Sniff-coding and inhibited neuronal responses in mice transduced with synapsin GCaMP6**.

**Supplementary Movie 2: Transient and sustained RTN neuronal responses to hypercapnia in synapsin GCaMP6 tranfected mice**.

**Supplementary Movie 3: Graded RTN neuronal response to hypercapnia in synapsin GCaMP6 tranfected mice**.

**Supplementary Movie 4: Rebound RTN neuronal response to hypercapnia in synapsin GCaMP6 tranfected mice**.

**Supplementary Movie 5: Transient and graded Raphe neuronal responses to hypercapnia in synapsin GCaMP6 tranfected mice**.

**Supplementary Movie 6: Transient Raphe neuronal response to hypercapnia in SERT GCaMP6 tranfected mice**.

**Supplementary Movie 7: Graded Raphe neuronal response to hypercapnia in SERT GCaMP6 tranfected mice**.

**Supplementary Movie 8: Inhibited Raphe neurons due to hypercapnia in SERT GCaMP6 tranfected mice**.

**Supplementary Movie 9: Transient and graded RTN glial responses to hypercapnia in GFAP GCaMP6 tranfected mice**.

**Supplementary Movie 10: Transient and graded Raphe glial responses to hypercapnia in GFAP GCaMP6 tranfected mice**.

## References

1 Guyenet, P. G., Stornetta, R. L. & Bayliss, D. A. Central respiratory chemoreception. The Journal of comparative neurology 518, 3883–3906, doi: 10.1002/cne.22435 (2010).

2 Nattie, E. Julius H. Comroe, Jr., Distinguished Lecture: Central chemoreception: then … and now. Journal of Applied Physiology 110, 1–8, doi: 10.1152/japplphysiol.01061.2010 (2011).

3 Huckstepp, R. T. & Dale, N. Redefining the components of central CO2 chemosensitivity--towards a better understanding of mechanism. J Physiol 589, 5561–5579, doi: 10.1113/jphysiol.2011.214759 (2011).

4 Smith, J. C., Morrison, D. E., Ellenberger, H. H., Otto, M. R. & Feldman, J. L. Brainstem projections to the major respiratory neuron populations in the medulla of the cat. J Comp Neurol 281, 69–96, doi: 10.1002/cne.902810107 (1989).

5 Nattie, E. E. & Li, A. Retrotrapezoid nucleus lesions decrease phrenic activity and CO2 sensitivity in rats. Respir Physiol 97, 63–77 (1994).

6 Mulkey, D. K. et al. Respiratory control by ventral surface chemoreceptor neurons in rats. Nat Neurosci 7, 1360–1369 (2004).

7 Ramanantsoa, N. et al. Breathing without CO2 Chemosensitivity in Conditional Phox2b Mutants. The Journal of Neuroscience 31, 12880–12888, doi: 10.1523/jneurosci.1721-11.2011 (2011).

8 Kumar, N. N. et al. Regulation of breathing by CO(2) requires the proton-activated receptor GPR4 in retrotrapezoid nucleus neurons. Science 348, 1255–1260, doi: 10.1126/science.aaa0922 (2015).

9 Richerson, G. B., Wang, W., Tiwari, J. & Bradley, S. R. Chemosensitivity of serotonergic neurons in the rostral ventral medulla. Respir Physiol 129, 175–189, doi: S0034568701002894 [pii] (2001).

10 Richerson, G. B. Serotonergic neurons as carbon dioxide sensors that maintain pH homeostasis. Nat Rev Neurosci 5, 449–461, doi: 10.1038/nrn1409 nrn1409 [pii] (2004).

11 Corcoran, A. E. et al. Medullary serotonin neurons and central CO(2) chemoreception. Respir Physiol Neurobiol, doi:S1569-9048(09)00100-1 [pii] 10.1016/j.resp.2009.04.014 [doi] (2009).

12 Ray, R. S. et al. Impaired respiratory and body temperature control upon acute serotonergic neuron inhibition. Science 333, 637–642, doi: 10.1126/science.1205295 (2011).

13 Pagliardini, S. et al. Active expiration induced by excitation of ventral medulla in adult anesthetized rats. J Neurosci 31, 2895–2905, doi: 10.1523/jneurosci.5338-10.2011 (2011).

14 Huckstepp, R. T., Cardoza, K. P., Henderson, L. E. & Feldman, J. L. Role of parafacial nuclei in control of breathing in adult rats. J Neurosci 35, 1052–1067, doi: 10.1523/jneurosci.2953-14.2015 (2015).

15 Huckstepp, R. T., Henderson, L. E., Cardoza, K. P. & Feldman, J. L. Interactions between respiratory oscillators in adult rats. eLife 5, doi: 10.7554/eLife.14203 (2016).

16 Ziv, Y. & Ghosh, K. K. Miniature microscopes for large-scale imaging of neuronal activity in freely behaving rodents. Curr Opin Neurobiol 32, 141–147, doi: 10.1016/j.conb.2015.04.001 (2015).

17 Jennings, J. H. et al. Visualizing hypothalamic network dynamics for appetitive and consummatory behaviors. Cell 160, 516–527, doi: 10.1016/j.cell.2014.12.026 (2015).

18 Resendez, S. L. et al. Visualization of cortical, subcortical and deep brain neural circuit dynamics during naturalistic mammalian behavior with head-mounted microscopes and chronically implanted lenses. Nat Protoc 11, 566–597, doi: 10.1038/nprot.2016.021 (2016).

19 Jakus, J., Halasova, E., Poliacek, I., Tomori, Z. & Stransky, A. Brainstem areas involved in the aspiration reflex: c-Fos study in anesthetized cats. Physiological research 53, 703–717 (2004).

20 Huckstepp, R. T. R., Cardoza, K. P., Henderson, L. E. & Feldman, J. L. Distinct parafacial regions in control of breathing in adult rats. PloS one 13, e0201485, doi: 10.1371/journal.pone.0201485 (2018).

21 Shi, Y. et al. Neuromedin B Expression Defines the Mouse Retrotrapezoid Nucleus. The Journal of Neuroscience 37, 11744, doi: 10.1523/JNEUROSCI.2055-17.2017 (2017).

22 Mitchell, R. A., Loeschcke, H. H., Massion, W. H. & Severinghaus, J. W. Respiratory responses mediated through superficial chemosensitive areas on the medulla Journal of Applied Physiology 18, 523–533 (1963).

23 Loeschcke, H. H. Central chemosensitivity and the reaction theory. J Physiol 332, 1–24 (1982).

24 Shams, H. Differential effects of CO2 and H+ as central stimuli of respiration in the cat. J Appl Physiol 58, 357–364 (1985).

25 Nattie, E. CO2, brainstem chemoreceptors and breathing. Prog Neurobiol 59, 299–331. (1999).

26 Nattie, E. E. Central chemosensitivity, sleep, and wakefulness. Respir Physiol 129, 257–268, doi: S003456870100295X [pii] (2001).

27 Milsom, W. K. Phylogeny of CO2/H+ chemoreception in vertebrates. Respir Physiol Neurobiol 131, 29–41 (2002).

28 Gourine, A. V., Llaudet, E., Dale, N. & Spyer, K. M. ATP is a mediator of chemosensory transduction in the central nervous system. Nature 436, 108–111, doi: nature03690 [pii] 10.1038/nature03690 (2005).

29 Gourine, A. V. et al. Astrocytes control breathing through pH-dependent release of ATP. Science 329, 571–575, doi: science.1190721 [pii] 10.1126/science.1190721 (2010).

30 Huckstepp, R. T. et al. Connexin hemichannel-mediated CO2-dependent release of ATP in the medulla oblongata contributes to central respiratory chemosensitivity. J Physiol 588, 3901–3920, doi: 10.1113/jphysiol.2010.192088 (2010).

31 Calizo, L. H. et al. Raphe serotonin neurons are not homogenous: electrophysiological, morphological and neurochemical evidence. Neuropharmacology 61, 524–543, doi: 10.1016/j.neuropharm.2011.04.008 (2011).

32 Iceman, K. E., Corcoran, A. E., Taylor, B. E. & Harris, M. B. CO2-inhibited neurons in the medullary raphe are GABAergic. Respir Physiol Neurobiol 203, 28–34, doi: 10.1016/j.resp.2014.07.016 (2014).

33 Iceman, K. E. & Harris, M. B. A group of non-serotonergic cells is CO2-stimulated in the medullary raphe. Neuroscience 259, 203–213, doi: 10.1016/j.neuroscience.2013.11.060 (2014).

34 Hennessy, M. L. et al. Activity of Tachykinin1-Expressing Pet1 Raphe Neurons Modulates the Respiratory Chemoreflex. The Journal of Neuroscience 37, 1807, doi: 10.1523/JNEUROSCI.2316-16.2016 (2017).

35 Andrews, C. G. & Pagliardini, S. Expiratory activation of abdominal muscle is associated with improved respiratory stability and an increase in minute ventilation in REM epochs of adult rats. Journal of Applied Physiology 119, 968–974, doi: 10.1152/japplphysiol.00420.2015 (2015).

36 Iizuka, M. & Fregosi, R. F. Influence of hypercapnic acidosis and hypoxia on abdominal expiratory nerve activity in the rat. Respir Physiol Neurobiol 157, 196–205, doi: 10.1016/j.resp.2007.01.004 (2007).

37 Marina, N. et al. Essential role of Phox2b-expressing ventrolateral brainstem neurons in the chemosensory control of inspiration and expiration. J Neurosci 30, 12466–12473, doi: 30/37/12466 [pii] 10.1523/JNEUROSCI.3141-10.2010 (2010).

38 Wu, Y. et al. Chemosensitivity of Phox2b-expressing retrotrapezoid neurons is mediated in part by input from 5-HT neurons. J Physiol 597, 2741–2766, doi: 10.1113/jp277052 (2019).

39 Takakura, A. C. et al. Peripheral chemoreceptor inputs to retrotrapezoid nucleus (RTN) CO2-sensitive neurons in rats. J Physiol 572, 503–523 (2006).

40 Ramanantsoa, N. et al. Ventilatory response to hyperoxia in newborn mice heterozygous for the transcription factor Phox2b. Am J Physiol Regul Integr Comp Physiol 290, R1691–1696, doi: 00875.2005 [pii] 10.1152/ajpregu.00875.2005 (2006).

41 Hosford, P. S. et al. CNS distribution, signalling properties and central effects of G-protein coupled receptor 4. Neuropharmacology 138, 381–392, doi: 10.1016/j.neuropharm.2018.06.007 (2018).

42 Bradley, S. R. et al. Chemosensitive serotonergic neurons are closely associated with large medullary arteries. Nat Neurosci 5, 401–402 (2002).

43 Johansen, S. L., Iceman, K. E., Iceman, C. R., Taylor, B. E. & Harris, M. B. Isoflurane causes concentration-dependent inhibition of medullary raphé 5-HT neurons <em>in situ</em>. Autonomic Neuroscience: Basic and Clinical 193, 51–56, doi: 10.1016/j.autneu.2015.07.002 (2015).

44 Brust, R. D., Corcoran, A. E., Richerson, G. B., Nattie, E. & Dymecki, S. M. Functional and developmental identification of a molecular subtype of brain serotonergic neuron specialized to regulate breathing dynamics. Cell Rep 9, 2152–2165, doi: 10.1016/j.celrep.2014.11.027 (2014).

45 Malheiros-Lima, M. R., Silva, J. N., Souza, F. C., Takakura, A. C. & Moreira, T. S. C1 neurons are part of the circuitry that recruits active expiration in response to the activation of peripheral chemoreceptors. eLife 9, doi: 10.7554/eLife.52572 (2020).

46 Silva, J. N., Oliveira, L. M., Souza, F. C., Moreira, T. S. & Takakura, A. C. Distinct pathways to the parafacial respiratory group to trigger active expiration in adult rats. American Journal of Physiology-Lung Cellular and Molecular Physiology 317, L402–L413, doi: 10.1152/ajplung.00467.2018 (2019).

47 Bagdy, E., Kiraly, I. & Harsing, L. G., Jr. Reciprocal innervation between serotonergic and GABAergic neurons in raphe nuclei of the rat. Neurochem Res 25, 1465–1473, doi: 10.1023/a:1007672008297 (2000).

48 Cao, Y., Matsuyama, K., Fujito, Y. & Aoki, M. Involvement of medullary GABAergic and serotonergic raphe neurons in respiratory control: Electrophysiological and immunohistochemical studies in rats. Neuroscience research 56, 322–331, doi: https://doi.org/10.1016/j.neures.2006.08.001 (2006).

49 McCall, R. B. GABA-mediated inhibition of sympathoexcitatory neurons by midline medullary stimulation. American Journal of Physiology-Regulatory, Integrative and Comparative Physiology 255, R605–R615, doi: 10.1152/ajpregu.1988.255.4.R605 (1988).

50 van de Wiel, J. et al. Connexin26 mediates CO2-dependent regulation of breathing via glial cells of the medulla oblongata. bioRxiv, 2020.2004.2016.042440, doi: 10.1101/2020.04.16.042440 (2020).

51 Rosin, D. L., Chang, D. A. & Guyenet, P. G. Afferent and efferent connections of the rat retrotrapezoid nucleus. J Comp Neurol 499, 64–89 (2006).

52 Ezure, K. & Manabe, M. Decrementing expiratory neurons of the Botzinger complex. II. Direct inhibitory synaptic linkage with ventral respiratory group neurons. Exp Brain Res 72, 159–166 (1988).

53 Richter, D. W. & Smith, J. C. Respiratory rhythm generation in vivo. Physiology (Bethesda) 29, 58–71, doi: 10.1152/physiol.00035.2013 (2014).

54 Yang, C. F., Kim, E. J., Callaway, E. M. & Feldman, J. L. Monosynaptic projections to excitatory and inhibitory preBötzinger Complex neurons. bioRxiv, 694711, doi: 10.1101/694711 (2019).

55 Ashhad, S. & Feldman, J. L. Emergent Elements of Inspiratory Rhythmogenesis: Network Synchronization and Synchrony Propagation. Neuron, doi: 10.1016/j.neuron.2020.02.005.

56 Mainland, J. & Sobel, N. The Sniff Is Part of the Olfactory Percept. Chemical senses 31, 181–196, doi: 10.1093/chemse/bjj012 (2005).

57 Simonyan, K., Saad, Z. S., Loucks, T. M. J., Poletto, C. J. & Ludlow, C. L. Functional neuroanatomy of human voluntary cough and sniff production. NeuroImage 37, 401–409, doi: https://doi.org/10.1016/j.neuroimage.2007.05.021 (2007).

58 Sobel, N. et al. Sniffing and smelling: separate subsystems in the human olfactory cortex. Nature 392, 282–286, doi: 10.1038/32654 (1998).

59 Macrides, F. Temporal relationships between hippocampal slow waves and exploratory sniffing in hamsters. Behavioral Biology 14, 295–308, doi: https://doi.org/10.1016/S0091-6773(75)90419-8 (1975).

60 Vanderwolf, C. H. The hippocampus as an olfacto-motor mechanism: were the classical anatomists right after all? Behav Brain Res 127, 25–47, doi: https://doi.org/10.1016/S0166-4328(01)00354-0 (2001).

61 Sobel, N. et al. Odorant-Induced and Sniff-Induced Activation in the Cerebellum of the Human. The Journal of Neuroscience 18, 8990, doi: 10.1523/JNEUROSCI.18-21-08990.1998 (1998).

62 Kurnikova, A., Deschênes, M. & Kleinfeld, D. Functional brain stem circuits for control of nose motion. Journal of neurophysiology 121, 205–217, doi: 10.1152/jn.00608.2018 (2018).

63 Deschenes, M., Kurnikova, A., Elbaz, M. & Kleinfeld, D. Circuits in the Ventral Medulla That Phase-Lock Motoneurons for Coordinated Sniffing and Whisking. Neural Plast 2016, 7493048, doi: 10.1155/2016/7493048 (2016).

64 Zoccal, D. B. et al. Interaction between the retrotrapezoid nucleus and the parafacial respiratory group to regulate active expiration and sympathetic activity in rats. Am J Physiol Lung Cell Mol Physiol 315, L891–L909, doi: 10.1152/ajplung.00011.2018 (2018).

65 Sreenivasan, V., Karmakar, K., Rijli, F. M. & Petersen, C. C. Parallel pathways from motor and somatosensory cortex for controlling whisker movements in mice. Eur J Neurosci 41, 354–367, doi: 10.1111/ejn.12800 (2015).

66 Huckstepp, R. T., Eason, R., Sachdev, A. & Dale, N. CO2-dependent opening of connexin 26 and related beta connexins. J Physiol 588, 3921–3931, doi: 10.1113/jphysiol.2010.192096 (2010).

67 Meigh, L. et al. CO2 directly modulates connexin 26 by formation of carbamate bridges between subunits. eLife 2, e01213, doi: 10.7554/eLife.01213 (2013).

68 Leirao, I. P., Silva, C. A., Jr., Gargaglioni, L. H. & da Silva, G. S. F. Hypercapnia-induced active expiration increases in sleep and enhances ventilation in unanaesthetized rats. J Physiol 596, 3271–3283, doi: 10.1113/JP274726 (2018).

