## Supplementary figures for "Analyzing the neuroglial brainstem circuits for respiratory chemosensitivity in freely moving mice"

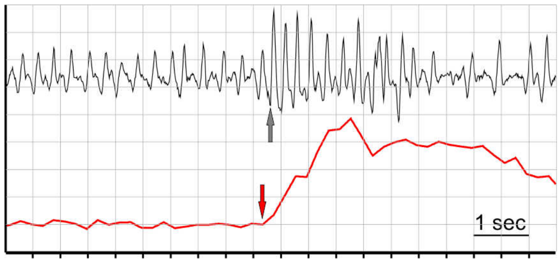


**Supplementary Figure 1: The onset of the RTN neuronal GCaMP signal relative to the start of sniff activity.** Red- calcium transient; gray- breathing changes.

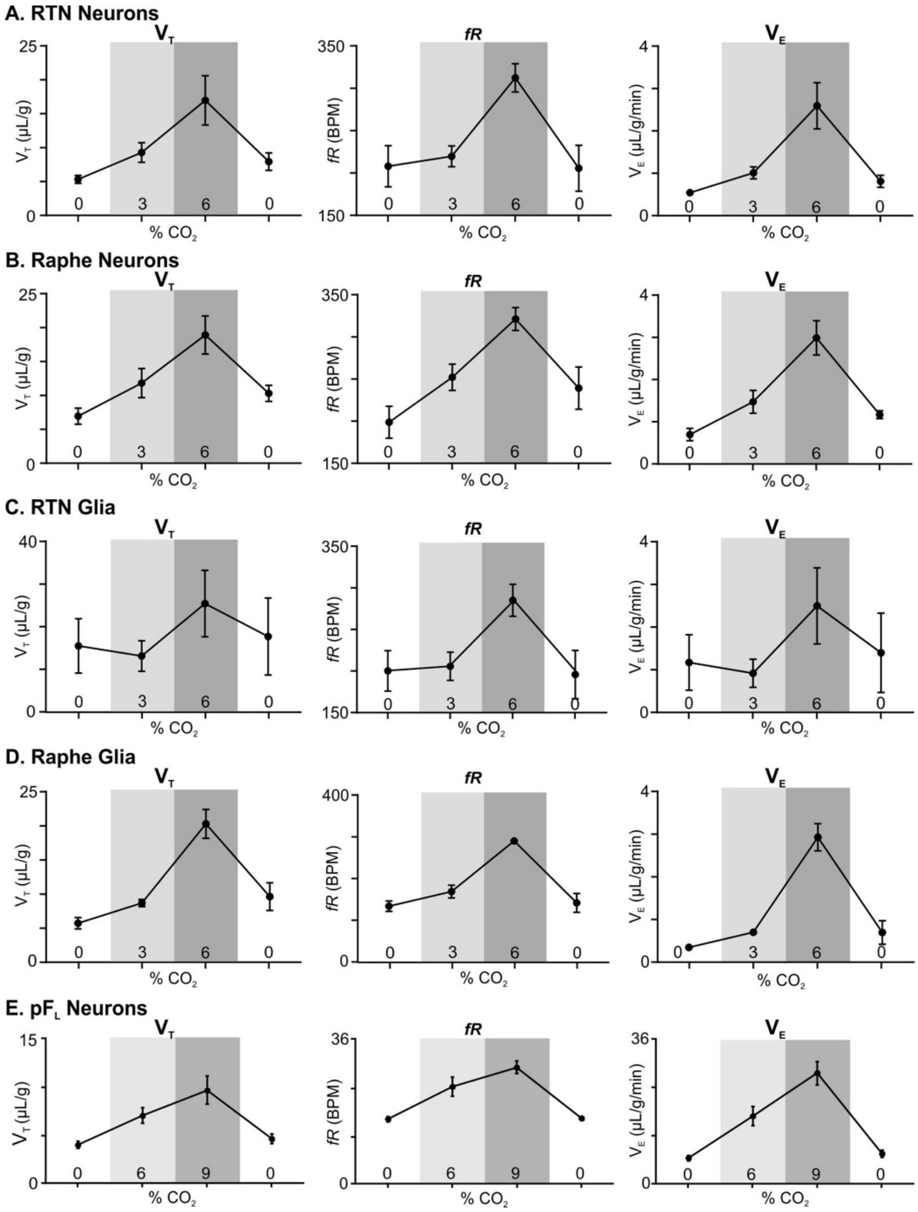


**Supplementary Figure 2: Change in breathing frequency (*f*R), tidal volume ($V_T$) and minute ventilation ($V_E$) in response hypercapnia in mice.** Change in tidal volume ($V_T$; left) breathing frequency (*f*R, bpm; centre), and minute ventilation ($V_E$; right) in response to changes in $CO_2$ concentration in (A) RTN neurons, (B) Raphe neurons, (C) RTN glia, (D) Raphe glia and (E) $pF_L$ neurons recorded mice.

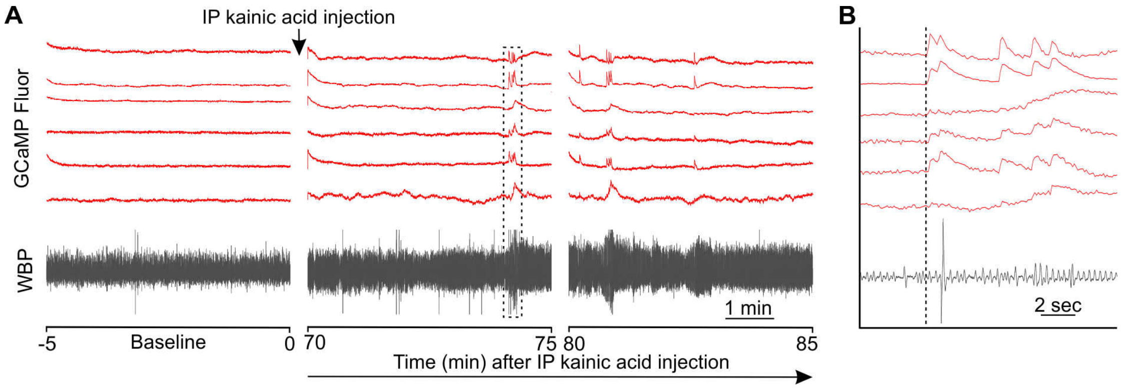


**Supplementary Figure 3: Seizure-induced increase in the activity of RTN neurons corelated with changes in breathing in anesthesized mice.** (A) Changes in mouse WBP before and after induction of seizures with intraperitoneal injection of kainic acid time matched with the activity of RTN neurons (red). The dotted square is changes in activity of neurons (ROIs) time matched with WBP and expanded in panel B. (B) Increase in the activity of RTN neurons correlated with changes in breathing. The start of the GCaMP6 transient is shown by the dotted line.

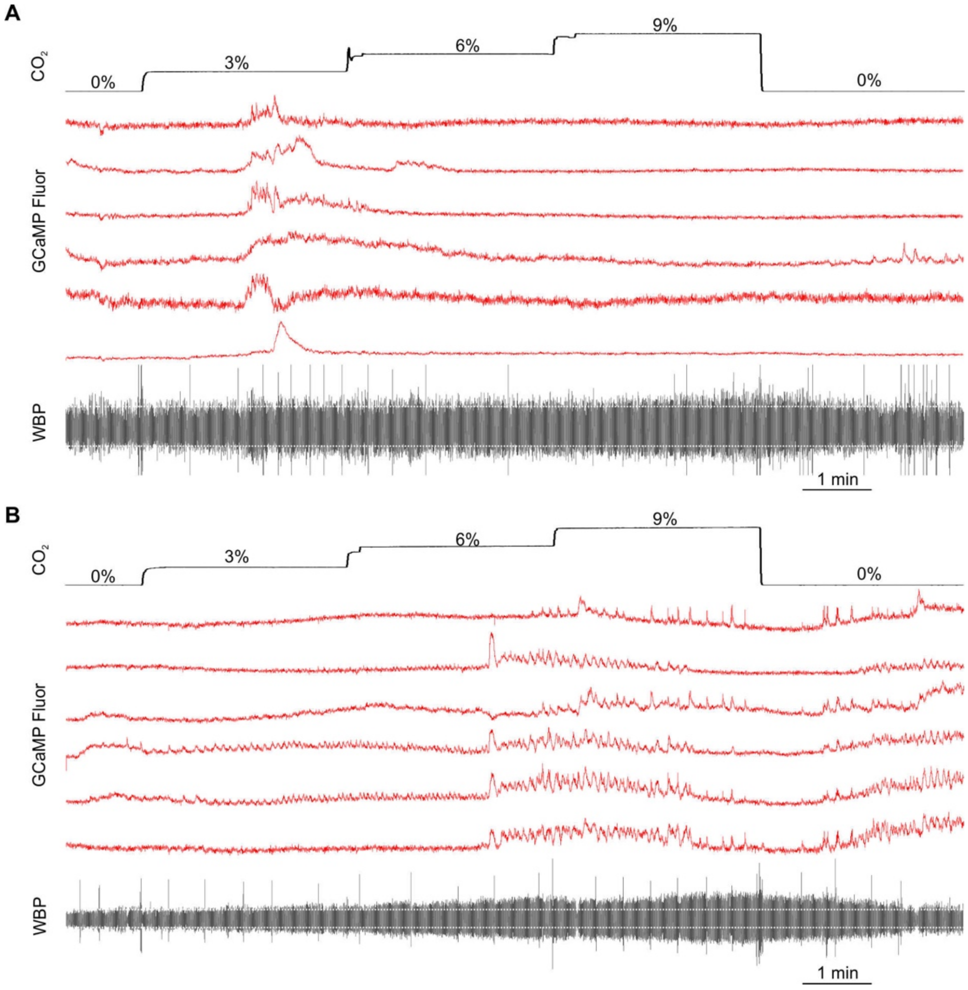


**Supplementary Figure 4: Hypercapnia induced activated RTN neurons in anesthesized mice.** (A-B) Mice under anesthesia evoked increased activity of RTN neuronal responses (red) to 3, 6 and 9% hypercapnia time-matched with WBP and $CO_2$ changes.
